# RNF40-dependent epigenetic regulation of actin cytoskeletal dynamics is required for HER2-driven mammary tumorigenesis

**DOI:** 10.1101/2020.02.24.962902

**Authors:** Florian Wegwitz, Evangelos Prokakis, Anastasija Pejkovska, Robyn Laura Kosinsky, Markus Glatzel, Klaus Pantel, Harriet Wikman, Steven A. Johnsen

**Author notes:** These authors contributed equally to this work (Florian Wegwitz, Evangelos Prokakis). **CORRESPONDING AUTHOR INFORMATION:** Steven A. Johnsen, Ph.D., Division of Gastroenterology and Hepatology, Mayo Clinic, 200 First St SW, Rochester, MN 55902.

## Abstract

The HER2-driven breast cancer subtype displays a particularly aggressive behavior. Alterations of the epigenome are common in cancers and represent attractive novel molecular therapeutic targets. Monoubiquitination of histone 2B (H2Bub1) by its obligate heterodimeric E3 ubiquitin ligase complex RNF20/RNF40 has been described to have tumor suppressor functions and loss of H2Bub1 has been associated with cancer progression. In this study, we utilized human tumor samples, cell culture models, and a mammary carcinoma mouse model with tissue-specific *Rnf40* deletion and identified an unexpected tumor-supportive role of RNF40 in HER2-positive breast cancer. We demonstrate that RNF40-driven H2B monoubiquitination is essential for transcriptional activation of RHO/ROCK/LIMK pathway components and proper actin cytoskeleton dynamics through a trans-histone crosstalk with histone 3 lysine 4 trimethylation (H3K4me3). Collectively, this work demonstrates a previously unknown essential role of RNF40 in HER2-positive breast cancer, revealing the RNF20/RNF40/H2Bub1 axis as a possible tumor context-dependent therapeutic target in breast cancer.

**Statement of significance:** HER2-positive breast cancer patients frequently develop resistance to anti-HER2 therapies. Here we demonstrate that RNF20/RNF40-mediated H2B monoubiquitination supports the oncogenic properties of cancer cells of this subtype by regulating actin dynamics. The RNF20/RNF40/H2Bub1 axis may therefore represent an attractive drug target for novel therapies.

## Introduction

Breast cancer (BC) is the most common form of cancer in the female population (1). The survival rates of BC vary greatly and strongly depend upon both early detection as well as the molecular subtype (2). Breast cancer can be separated into at least four distinct molecular subtypes based on the expression of the estrogen receptor (ER), progesterone receptor (PR) or human epidermal growth factor receptor 2 (HER2) receptor. Notably, the HER2-positive and triple negative (HER2-, ER-, PR-) BC subtypes are generally more invasive and display a poorer prognosis compared to hormone receptor-positive (ER+ or/and PR+) BC (3). Importantly, while current anti-HER2 therapies are initially highly effective for many BC patients with HER2-positive tumors, a significant number of patients develop tumors refractory to therapy and display tumor relapse and disease progression (4). Thus, new approaches are necessary to combat HER2-positive breast cancer.

Precision oncology approaches aim to utilize or develop novel targeted therapies which exploit tumor-specific dependencies and/or vulnerabilities based on specific molecular alterations present in a given tumor or molecular subtype (5). In addition to genetic alterations that occur in cancer, a large body of emerging evidence has demonstrated the additional importance of epigenetic alterations in tumorigenesis and tumor progression. These alterations can occur either through the direct mutation of genes encoding epigenetic regulatory proteins or as secondary events downstream of signaling pathways altered as a result of other genetic changes (6). These changes can result in altered patterns of DNA methylation, post-translational histone modifications, and changes in chromatin accessibility or chromatin architecture. Due to the reversible nature of many of these changes, numerous substances are currently in various stages of pre-clinical and clinical testing to determine their efficacy as anti-cancer therapies (7).

Previous work from our lab and others revealed a particular importance of histone 2B monoubiquitination (H2Bub1) in controlling cellular differentiation (8–10) and demonstrated that H2Bub1 levels are decreased in ER-positive BC compared to normal adjacent epithelium (11,12). These findings have led to the hypothesis that H2Bub1, catalyzed by the obligate heterodimeric Ring Finger Protein 20 and 40 (RNF20/RNF40) E3 ubiquitin ligase complex, has a tumor suppressive function. This hypothesis has been further supported by studies investigating the function of RNF20 and RNF40 (13–17). In contrast, we and others have uncovered tumor supportive roles of RNF20 and RNF40 in colorectal cancer (18,19) and androgen-dependent prostate cancer (20). Therefore, together these findings suggest that RNF20/RNF40-driven H2B monoubiquitination plays a context-dependent role in cancer.

At the molecular level, H2Bub1 is localized across the body of active genes (21) and is closely coupled to transcriptional elongation (22–25). Studies in both yeast and human cells have revealed a particular coupling of H2Bub1 and the trimethylation of lysines 4 and 79 of histone 3 (H3K4me3 and H3K79me3, respectively) near the transcriptional start site and transcribed regions, respectively, of active genes (22,26–30). Past studies have reported an extension of H3K4me3 into the transcribed region of genes displaying a particularly high transcriptional elongation rate (31), as well as a close link between H2Bub1 and transcriptional elongation (23). Consistently, we recently demonstrated that loss of RNF40-mediated H2B monoubiquitination results in the narrowing of H3K4me3 domains near the transcriptional start site (TSS) of important cell fate-determining genes displaying high elongation rates (22).

In this study we sought to examine the role of RNF40-mediated H2B monoubiquitination in the HER2-driven subtype of BC. Our studies using a tissue-specific transgenic and gene ablation approach demonstrate for the first time that RNF40 exerts a profound tumor supportive function in the biology of HER2-driven mammary carcinoma. In support of these *in vivo* findings, we show that RNF40 silencing leads to decreased cell proliferation and specific transcriptional and epigenetic changes in HER2-positive human BC cells lines. Finally, we unveil a previously undescribed role of RNF40-mediated H2B monoubiquitination in driving the expression of specific genes regulating actin cytoskeleton dynamics (*in vitro* and *in vivo*) and the activity of the downstream FAK-driven signaling cascade.

## RESULTS

### RNF40 is highly expressed in HER2-positive BC

While we and others have uncovered potential differing tumor-supportive or tumor-suppressive roles of H2Bub1 and its E3 ligases RNF20 and RNF40 in ER-positive and triple negative BC (11,12,16), the role of this epigenetic pathway in HER2-positive BC is currently unclear. Therefore, we investigated RNF40 expression and H2Bub1 levels by immunohistochemical staining of 176 primary BC tumors and 78 brain metastases. Interestingly, examination of RNF40 and H2Bub1 staining revealed that all analyzed HER2-positive BC samples were positive for both markers (Fig. 1A-C and Fig S1A). Moreover, HER2-positive metastatic BC samples showed a particularly high expression of RNF40 compared to primary tumors (Fig. 1A-C and Fig S1A). We next examined the relationship between *RNF40* mRNA levels and survival in HER2-positive BC patients using publically available data and observed that high levels of *RNF40* expression were associated with a reduced overall and relapse-free survival (Fig. S1C and Fig. 1D). Strikingly, in the same dataset, RNF40 mRNA expression was found to be significantly higher in HER2-positive breast cancer tissues compared to normal mammary tissues (Fig. S1B). In summary, these data suggest a potentially unexpected tumor supportive role of RNF40 in HER2-positive BC.

**Fig. 1:**
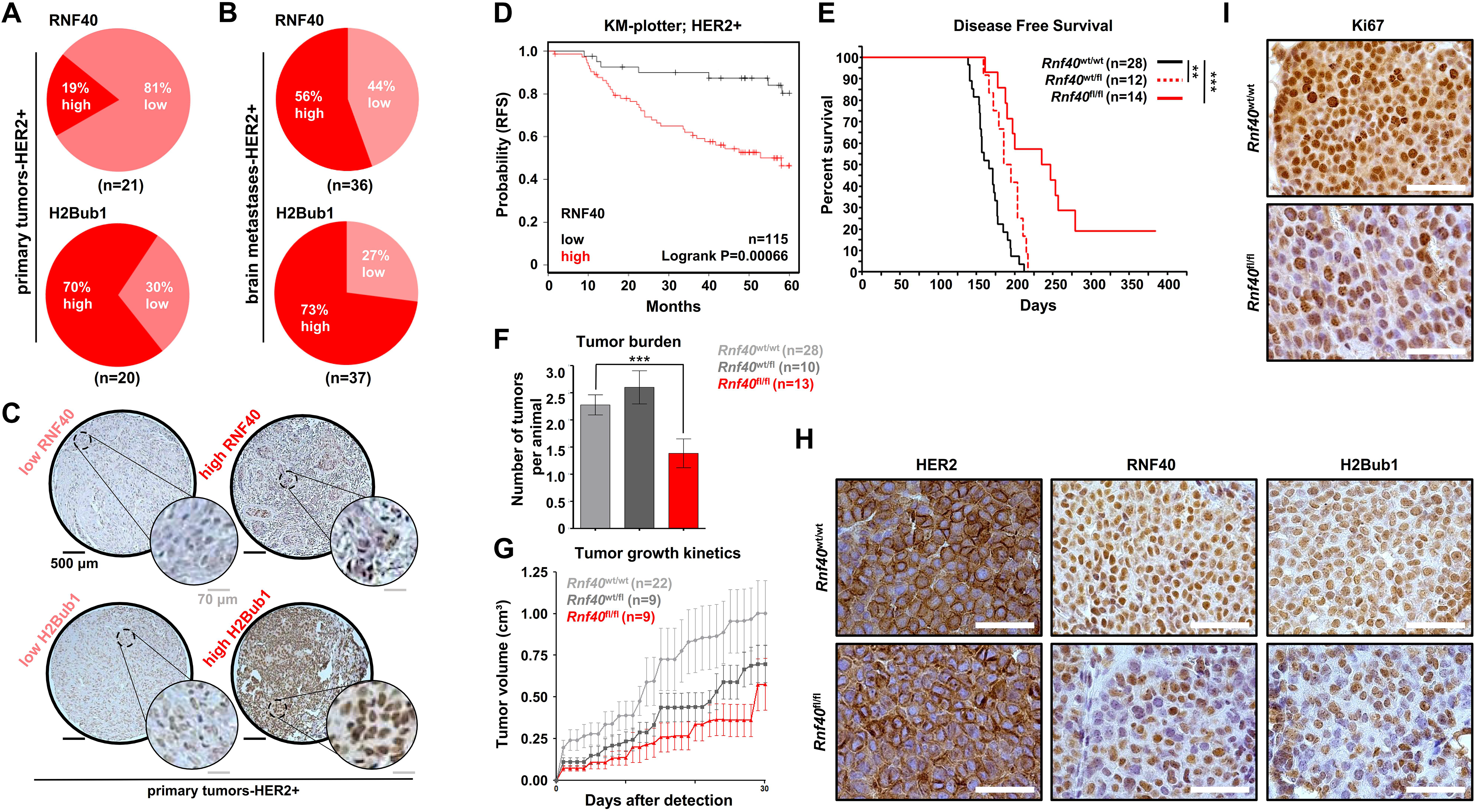
RNF40 and H2Bub1 are maintained in HER2-positive breast cancer. **A-B:** TMAs with primary mammary carcinoma (n=178) and brain metastasis (n=78) lesions were stained for RNF40 and H2Bub1 by IHC. Distribution of RNF40 and H2Bub1 staining intensity in all BCs (left panels) and in the HER2-positive subtype (right panel) in primary tumors **(A)** and brain metastases **(B)**. **C)** Representative pictures of low and high H2Bub1 and RNF40 staining intensity in primary HER2-positive BC specimens. **D)** Relapse-free survival plot (RFS) of HER2-positive breast cancer patients with low and high gene expression of *RNF40*, using the online tool of kmplot.com. **E)** Disease-free survival of *Rnf40*^wt/wt^ compared to *Rnf40*^wt/fl^ or *Rnf40*^fl/fl^ mice. Logrank test. **F)** Bar graph depicting the average number of observed tumors per animal in each transgenic mouse cohort. Student t-test. **G)** Tumor growth kinetics of all transgenic mouse cohorts. **H)** Representative images of immunohistochemical staining of RNF40, H2Bub1 and HER2 in the *Rnf40*^wt/wt^ and *Rnf40*^fl/fl^ mammary carcinomas. Scale bar (white): 100 μm **I)** Immunohistochemical detection of the Ki67 proliferation marker. Scale bars (white): 100 μm. **p-val<0.01, ***p-val<0.005. Error bars: standard error of the mean (SEM).

### RNF40 plays a tumor supportive function in *Erbb2*-driven mammary carcinoma *in vivo*

Since RNF40 expression and activity were largely maintained in human HER2-positive BC, we hypothesized that a loss of RNF40 may impair HER2-driven tumorigenesis in a mouse model system. Therefore, to test this hypothesis, we utilized the MMTV-*Erbb2* genetic mouse model initiating HER2-positive mammary lesions upon overexpression of the *Erbb2* proto-oncogene (coding for HER2) specifically in mammary epithelial cells (32). We generated a tri-transgenic MMTV-*Erbb2*; MMTV-Cre; *Rnf40*^*f*lox^ mouse line with mammary tissue-specific co-expression of HER2 and Cre-recombinase, and a floxed *Rnf40* allele. This approach enabled us to achieve a simultaneous HER2 overexpression and mammary epithelium-specific ablation of *Rnf40* (18,22). Consistent with our findings in human HER2-positive BC lesions, MMTV-*Erbb2; Rnf40*^wt/wt^ tumors did not display a loss of either RNF40 or H2Bub1 (Fig. 1H) when compared to the adjacent normal mammary epithelium (Fig. S1F). Moreover, immunohistochemical analyses confirmed that HER2 expression was unaffected by *Rnf40* deletion (Fig. 1H). However, both heterozygous (*Rnf40*^wt/fl^), and especially homozygous loss of *Rnf40* (*Rnf40*^fl/fl^) resulted in dramatically increased tumor-free survival of MMTV-*Erbb2* animals (Fig. 1E). Remarkably, despite the high tumor incidence in this mouse model (100% of *Rnf40*^wt/wt^ mice developed tumors after 220 days), 2 out of 14 *Rnf40*^fl/fl^ animals (14%) never developed tumors even after 18 months observation (Fig. 1E). Analyses of the tumor burden revealed that *Rnf40*^fl/fl^ mice developed significantly fewer tumors than *Rnf40*^wt/wt^ (Fig. 1F) and loss of *Rnf40* led to strongly reduced tumor growth kinetics (Fig. 1G). Notably, *Rnf40* loss did not induce morphological changes, as visible in H&E staining of the *Rnf40*^wt/wt^ and *Rnf40*^fl/fl^ tumors (Fig. S1D). To estimate the efficiency of *Rnf40* deletion in this model, we performed RNF40 immunohistochemical staining in *Rnf40*^wt/wt^ and *Rnf40*^fl/fl^ tumors. Consistent with the lack of a complete block in tumor incidence and growth, *Rnf40*^fl/fl^ lesions displayed a heterogeneous pattern of RNF40 expression (Fig. 1H), suggesting that the few tumors that did develop in this model were largely caused by an incomplete loss of the *Rnf40* allele. This is further supported by the observation that both H2Bub1, as well the proliferation marker Ki67, displayed a similar heterogeneous expression pattern as RNF40 (Fig. 1H-I and Fig. S1E). Similar effects have been reported in a number of other tumor types and with various Cre models, where occasional tumors did appear, which all retained some expression of the floxed essential tumor driver gene of interest (33,34). Thus, we posit that these findings provide further support for the essential role of RNF40 in HER2-driven tumorigenesis to the extent that rare, RNF40/H2Bub1-expressing “escaper” cells are positively selected for during tumorigenesis and tumor progression. Taken together, these results demonstrate that RNF40 plays an essential supportive function in HER2-driven mammary tumor initiation and progression.

### RNF40 loss impairs oncogenic properties of HER2-positive BC cells *in vitro*

We next sought to investigate the underlying molecular mechanisms determining the dependence of HER2 positive BC on RNF40. In order to achieve this goal, we selected two different human HER2-positive BC cell lines (HCC1954, SKBR3) and assessed different parameters related to their tumorigenic properties following siRNA-mediated RNF40 knockdown. RNF40 depletion and concomitant loss of H2Bub1 in both cell lines (Fig. 2A, Fig. S2A-B) resulted in reduced cellular proliferation compared to control transfected cells (Fig. 2B-C; Fig. S2A-B). Furthermore, growth kinetics (Fig. 2C and Video in supplements), clonogenic capacity (Fig. 2D) and tumor sphere formation (Fig. 2E) were strongly impaired upon RNF40 loss in both cell lines. In support of these results, an analysis of the data in the DepMap portal (https://depmap.org/), which contains information for gene essentiality deriving from various RNAi and CRISPR screens, revealed that HER2-positive breast cancer cell lines are significantly impacted by RNF40 loss by CRISPR/Cas9-mediated deletion, further confirming a particularly important role for RNF40 in this subtype of breast cancer (Fig. S2C). Consistently, the levels of the proliferation marker Ki67 were strongly reduced in both HER2-positive BC cell lines upon RNF40 depletion (Fig. 2F). Finally, we also tested the migration potential of HCC1954 cells upon RNF40 depletion in trans-well migration (Fig. 2G) and gap closure assays (Fig. S2D). Notably, both approaches showed impaired cellular motility upon loss of RNF40. Together these findings support our in *vivo* findings that RNF40 expression is essential for maintaining tumorigenic properties of HER2-positive BC cells both *in vitro* and *in vivo*.

**Fig. 2:**
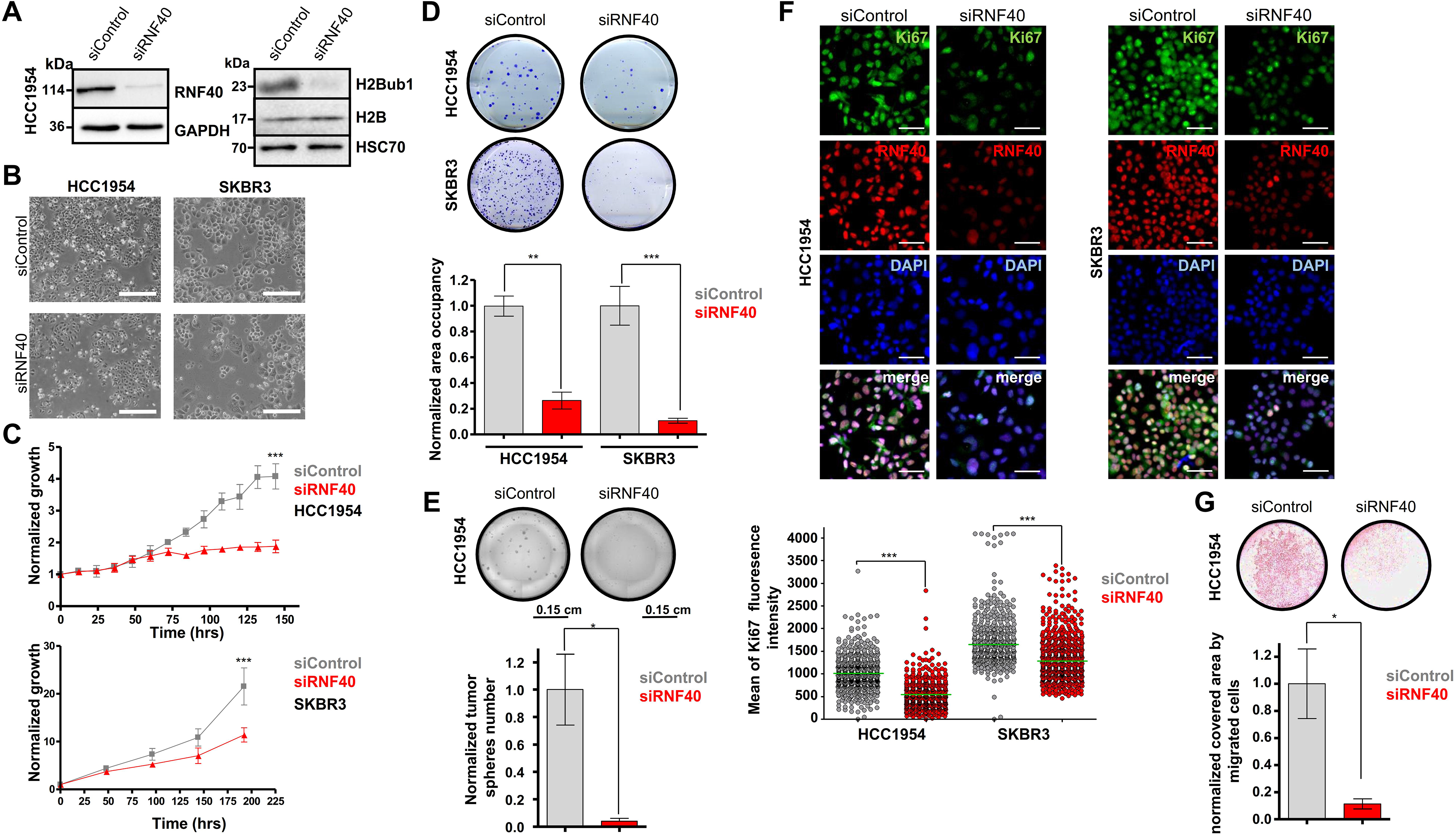
RNF40 loss impairs oncogenic properties of HER2-positive BC cells *in vitro*. **A)** Western blot validation of RNF40 knockdown efficiency and decreased H2Bub1 levels in HCC1954 cells. **B)** Representative bright-field pictures of control and RNF40 siRNA-transfected HCC1954 and SKBR3 cells. Scale bars (white): 500 μm. **B** and **C**: Proliferation curves **(C)** and clonogenic assays **(D)** of control and RNF40-depleted HCC1954 and SKBR3 cells. Quantification of the occupied area in clonogenic assays is shown for both cell lines (D, lower panel). Student t-test. **E)** Tumor sphere formation assay of control and RNF40-depleted HCC1954 cells (upper panel). Quantification of the respective tumor spheres number normalized to the control condition (lower panel). Student t-test. **F)** Representative pictures from immunofluorescence detection of RNF40 and the Ki67 proliferation marker in control and RNF40-depleted HCC1954 and SKBR3 cells. Scale bars (white) = 60 μm (upper panel). Quantification of the Ki67 immunofluorescence intensity of single nuclei in control and RNF40-depleted HCC1954 and SKBR3 cells (lower panel). The median intensity values of the respective groups are provided as green bars. Mann-Whitney test. **G)** Boyden-chamber-based migration assay of control and RNF40-depleted HCC1954 cells with representative results (upper panel) and the corresponding quantification (lower panel). *p-val<0.05, **p-val<0.01, ***p-val<0.005. Error bars of all quantification analyses: SEM.

### RNF40 regulates actin cytoskeleton-related genes in HER2-positive BC cells

Based on the dramatic effects observed on HER2-driven tumorigenesis *in vivo* and *in vitro*, we tested whether the activity of the signaling cascade downstream of HER2 may be directly affected by RNF40 loss. However, while the HER2 inhibitor Lapatinib (lap) significantly blocked ERK and AKT phosphorylation in the HER2-positive HCC1954 cell line, both pathways remained intact following RNF40 depletion (Fig. 3A). Therefore, given the direct epigenetic role of H2Bub1 in facilitating gene transcription, we performed mRNA-sequencing analyses of HCC1954 cells following RNF40 depletion or control siRNA-transfection and identified 360 up- and 324 down-regulated genes (| log2 fold change|>0.6; p-val<0.05) (Fig. 3B). Consistent with our previous findings in colorectal cancer (19), Gene Set Enrichment Analysis (GSEA) identified a significant enrichment for a gene signature associated with hallmarks of apoptosis, potentially explaining the reduced oncogenic properties (Fig. 3C and S3B). Increased apoptosis could be confirmed by microscopic time lapse analyses (see videos in supplements), higher levels of cleaved caspase 3 and cleaved PARP in Western blot (Fig. 3D) and an increase in Annexin V-positive cells in FACS-based analyses in HCC1954 cells (Fig. 3E). Given the fact that RNF40 depletion not only resulted in decreased cell number, but also dramatically affected cell migration, we performed additional gene ontology analyses using the EnrichR tool (https://amp.pharm.mssm.edu/Enrichr/) and identified an enrichment of genes associated with the actin cytoskeleton regulatory pathway as being downregulated following RNF40 depletion (Fig. 3F and S3C). We selected genes from this set and confirmed the downregulation of Vav Guanine Nucleotide Exchange Factor 3 (*VAV3*), Rho Associated Coiled-Coil Containing Protein Kinase 1 (*ROCK1*), LIM Domain Kinase 2 (*LIMK2*) and Profilin 2 (*PFN2*), which directly control filamentous actin dynamics, both at the mRNA (Fig. 3G and Fig. S3A) and protein (ROCK1, VAV3; Fig. 3H) levels.

**Fig. 3:**
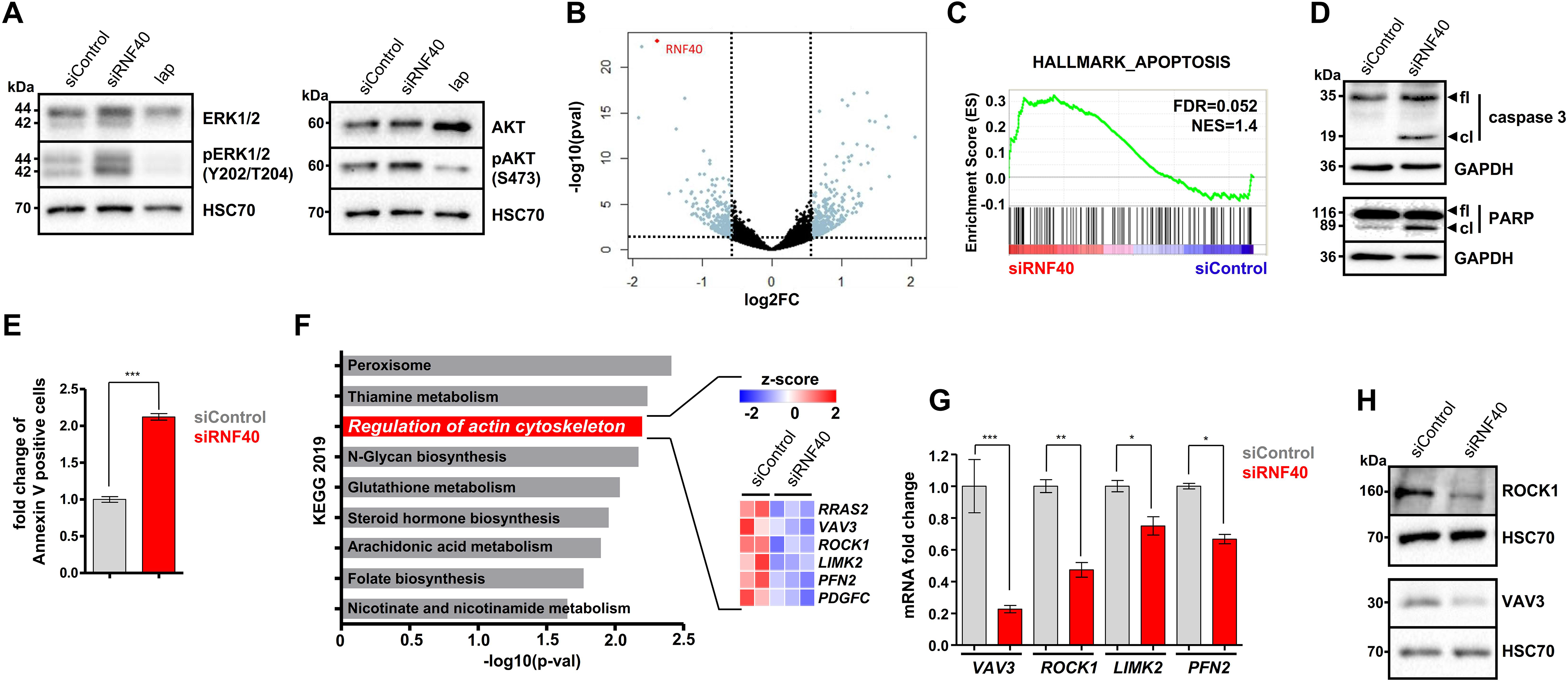
RNF40 loss increases apoptosis and impairs the expression of key components of the actin regulatory pathway in HER2-positive BC cells. **A)** Western blot analysis of the total and phosphorylated forms of ERK1/2 and AKT in control and RNF40-depleted HCC1954 cells. 1 μM Lapatinib (lap) was applied for 12 hours as a positive control. **B)** Volcano plot displaying gene expression changes occurring in HCC1954 cells upon RNF40 depletion and measured by mRNA sequencing. **C)** Gene Set Enrichment Analysis (GSEA) of the mRNA sequencing data significantly enriched for “Hallmark_Apoptosis” geneset enriched in the RNF40-depleted condition. **D)** Western blot analysis showing that the markers of apoptosis, the cleaved forms of caspase 3 and PARP, are at higher levels in RNF40-depleted HCC1954 cells compared to the control condition. **E)** Annexin V assay of control and RNF40-depleted HCC1954 cells. **F)** Pathway enrichment analysis (EnrichR web tool) showing that genes significantly downregulated upon RNF40 knockdown are enriched for the KEGG 2019 signature “Regulation of actin cytoskeleton”. A heatmap depicting the differential expression of genes involved in this signature is provided in the right panel). **G-H:** The identified signature was validated via qRT-PCR **(G)** and western blot **(H)** for selected genes in HCC1954 cells. Student t-test. *p-val<0.05, **p-val<0.01, ***p-val<0.005. Error bars of all quantification analyses: SEM.

Phosphorylation of the cofilin protein by LIMK downstream of ROCK1 plays an important role in controlling actin cytoskeleton dynamics (35). In its active unphosphorylated form, cofilin destabilizes F-actin and leads to actin depolymerization. We therefore assessed cofilin phosphorylation (p-cofilin) via western blotting and observed strongly reduced levels in RNF40-depleted HCC1954 cells (Fig. 4A). Consistently, phalloidin staining further confirmed the impairment of F-actin formation upon RNF40 depletion in HCC1954 (Fig. 4C) and SKBR3 cells (Fig. S4A) and these effects could be phenocopied by inhibition of ROCK1 by RKI-1447 (Fig. 4A and C) or siRNA-mediated knockdown of VAV3 (Fig. 4B). Importantly, these effects could also be confirmed *in vivo* where cofilin phosphorylation was also significantly decreased in *Rnf40*^fl/fl^ tumors compared to wild type *Rnf40* tumors (Fig. 4D). Together, these data confirm the *in vitro* and *in vivo* importance of RNF40 in controlling actin cytoskeletal dynamics in HER2-positive BC.

**Fig. 4:**
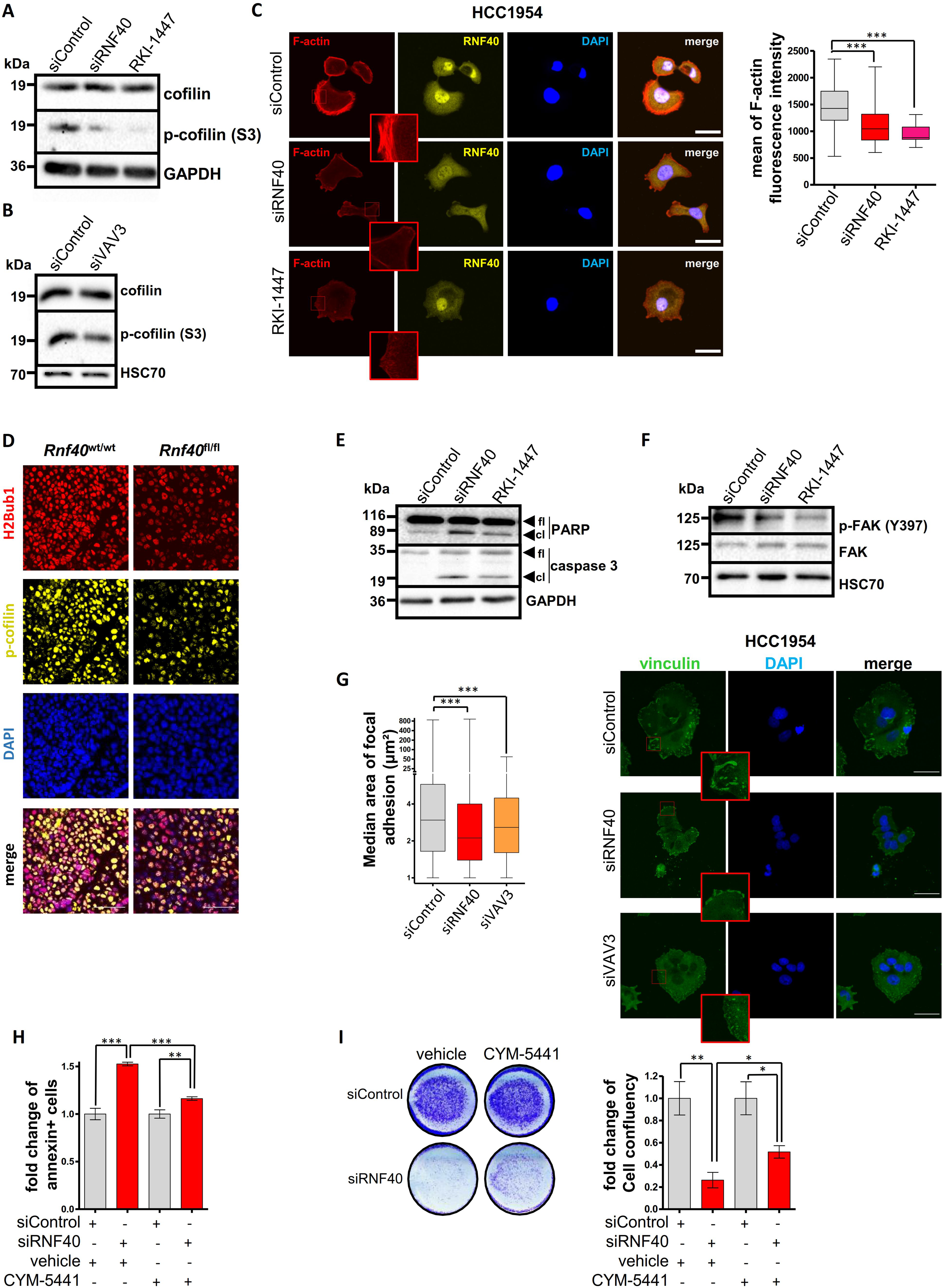
RNF40 controls the actin regulatory pathway to sustain the viability of HER2-positive BC cells *in vitro* and *in vivo*. **A)** Western blot analysis showing a reduction of phosphorylated cofilin (p-cofilin) upon RNF40 knockdown and ROCK inhibitor treatment (RKI-1447) in HCC1954 cells. **B)** Western blot analysis showing a reduction of phosphorylated cofilin in VAV3-depleted HCC1954 cells. **C)** Representative pictures of immunofluorescence staining for F-actin in control, RNF40-depleted and RKI-1447-treated (ROCK inhibitor) HCC1954 cells (right panel). Quantification of F-actin intensity in the respective conditions (left panel). Scale bars (white) = 50 μm. Mann-Whitney test. **D)** Representative pictures of p-cofilin detected by immunofluorescence in the murine *Rnf40*^wt/wt^ and *Rnf40*^fl/fl^ tumors. Western blot analysis assessing cleaved PARP and cleaved caspase 3 levels (cl: cleaved, fl: full length) **(E)** or phosphorylated and total FAK **(F)** in control, RNF40-depleted and RKI-1447-treated HCC1954 cells. **G)** Representative immunofluorescence pictures of vinculin in control, RNF40-depleted and VAV3-depleted HCC1954 cells (right panel). Bar graph displaying the median focal adhesion area in the respective conditions (left panel). Scale bars (white) = 50 μm. Mann-Whitney test. **H-I)** Annexin V assay **(H)** and proliferation assay **(I)** of control and RNF40-depleted HCC1954 cells with and without the S1PR agonist CYM-5441. Quantification of cell confluency (right panel). Student t-test. *p-val<0.05, **p-val<0.01, ***p-val<0.005. Error bars of all quantification analyses: SEM.

In addition to the established role of the ROCK1 pathway in controlling actin cytoskeletal dynamics, this pathway also plays a central role in suppressing apoptosis and potentiating cell survival (36–39). Notably, the potent ROCK inhibitor RKI-1447 was shown to elicit a pronounced anti-tumorigenic effect on the same *Erbb2*-driven mammary carcinoma mouse model used in our current study (37). Thus, we hypothesized that the dysregulation of the ROCK1-depedent actin regulatory pathway may play a central role in the apoptotic phenotype induced by RNF40 loss. Indeed, RKI-1447 treatment led to impaired HCC1954 cell proliferation (Fig. S4B) and the induction of caspase 3 cleavage (Fig. 4E). We therefore conclude that RNF40 has a decisive impact on the apoptotic rate of HER2-positive BC cells by regulating important members of the VAV3-ROCK1-LIMK2-PFN2 axis.

Focal adhesion complexes are cell-to-substrate adhesion structures that are tightly coupled to F-actin dynamics and significantly contribute to preserving anti-apoptotic pathways via the Focal Adhesion Kinase (FAK) (38,40). Since we demonstrated a crucial function of RNF40 in regulating the formation of F-actin stress fibers, we hypothesized that RNF40 depletion may influence the cell growth potential of HER2-positive BC cells by impairing focal adhesion signaling via impaired actin dynamics. To test this hypothesis, we first estimated the median area of focal adhesions via immunofluorescent staining for vinculin, one of the molecules which bridges focal adhesion complexes and F-actin. Indeed, the area of focal adhesion was substantially decreased upon RNF40 depletion and these effects could be phenocopied by VAV3 depletion (Fig. 4G). Furthermore, the levels of active phosphorylated FAK (p-FAK) were decreased upon RNF40 depletion in HCC1954 cells and could be phenocopied by ROCK inhibition (Fig. 4F). Moreover, consistent with these findings, direct inhibition of FAK led to a significant decrease in HCC1954 cell number (Fig. S4D).

To ensure the causality of the impaired tumorigenic phenotype of RNF40-silenced HCC1954 cells due to an impaired actin regulatory pathway, we examined the effects of restoring this signaling cascade. For this purpose, we treated HCC1954 cells with an allosteric sphingosine 1-phosphate receptor-3 agonist (CYM-5441), which was shown to activate actin polymerization as well as increase cancer stem cell expansion in BC (41,42). Treatment of RNF40-depleted HCC1954 cells with CYM-5441 significantly rescued apoptosis as measured by Annexin V staining (Fig. 4I) and caspase 3/7 activity (Fig. S4C). Additionally, treatment with either CYM-5441 (Fig. 4J) or lysophosphatidic acid (Fig. S4E), which has also been shown to activate this pathway (43), partially rescued the impaired proliferation of HCC1954 cells following RNF40 depletion. Notably, this rescue was prevented by treatment with RKI-1447 (Fig. S4F-G), confirming that partial restoration of the actin regulatory pathway is central to the observed rescuing effects. Collectively, these findings establish RNF40 as an important regulator of HER2-positive BC cell viability by regulating the actin regulatory process *in vitro* and *in vivo* via the regulation of the VAV3-ROCK1-LIMK2-PFN2 and focal adhesion signaling cascade.

### RNF40 regulates the VAV3-ROCK-LIMK2-PFN2 axis through H2Bub1-H3K4me3 trans-histone crosstalk

Previous studies from other groups demonstrated a crosstalk between H2Bub1 and H3K4 tri-methylation (H3K4me3) both in yeast and human systems (26,29,30). Moreover, we recently demonstrated that RNF40-mediated H2B monoubiquitination specifically governs the transcriptional start site- (TSS-) proximal broadening of H3K4me3 into the transcribed region to facilitate transcriptional elongation of a number of moderately H2Bub1-marked genes in mouse embryo fibroblasts (MEFs) (22). To examine if RNF40 controls the expression of genes of the actin regulatory network by modulating H2Bub1 and H3K4me3 levels, we performed chromatin immunoprecipitation sequencing (ChIP-seq) analyses for H2Bub1 and H3K4me3 in HCC1954 cells (Fig. S5A). Strikingly, consistent with our previous findings (22), RNF40-dependent genes showed lower levels of H2Bub1 occupancy compared to unregulated genes or genes up-regulated following RNF40 depletion (Fig. 5A). Given our previous finding that H3K4me3 “peak narrowing” is a distinct epigenetic feature involved in the regulation of RNF40-dependent genes, we identified peaks displaying either an increase or a global or partial (3’ narrowing) decrease of H3K4me3 occupancy upon RNF40 silencing in HCC1954 cells (Fig. 5B). We then utilized the identified regions for differential binding (Diffbind) analyses and observed that the majority of the regions influenced by RNF40 depletion markedly lost H3K4me3 (8,518 regions), whereas only a few regions gained H3K4me3 occupancy (351 regions) (Fig. 5C and Fig. 5D). Interestingly, most of the regions showing decreased H3K4me3 (Fig. 5C) were located proximal to TSS regions (Fig. S5B). Moreover, regions displaying no changes in H3K4me3 occupancy show only a mild peak narrowing, while regions displaying a significant loss of H3K4me3 occupancy exhibit a stronger peak narrowing upon RNF40 depletion (Fig. 5D and S5C). Importantly, consistent with our gene expression analyses, TSS-associated regions displaying decreased H3K4me3 occupancy following RNF40 depletion included genes associated with the actin regulatory pathway signature (Fig. S5D).

**Fig. 5:**
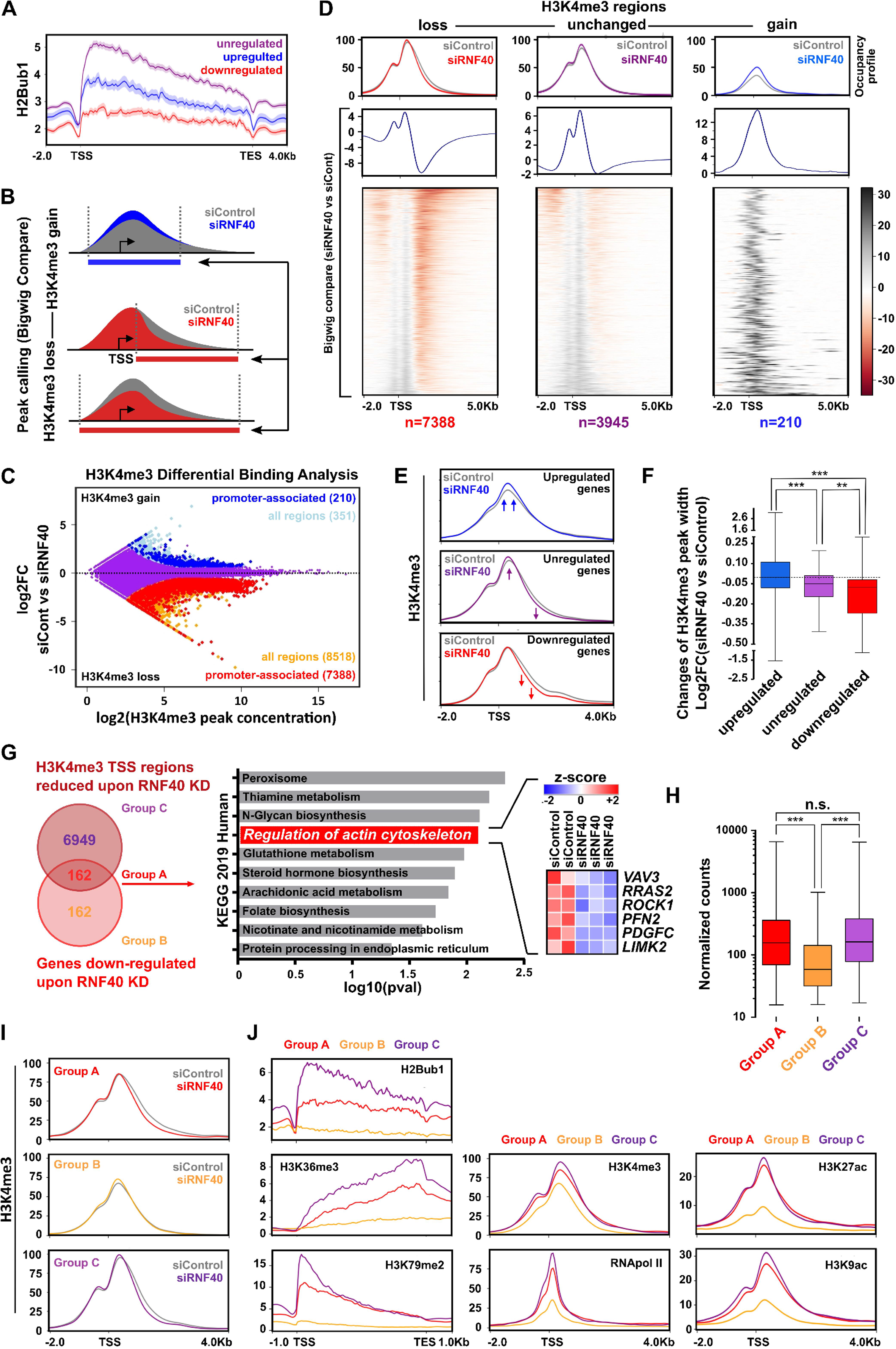
RNF40 regulates gene expression of important members of the RHO-ROCK axis in an H2Bub1/H3K4me3-dependent manner. **A)** Gene body H2Bub1 occupancy profiles on down-, up- and unregulated genes upon RNF40 depletion (regulated genes |log2FC|≥0.6, p-val<0.05, unregulated genes |log2FC|≤0.1, p-val>0.95). **B)** Schematic workflow showing the procedure utilized to identify regions losing or gaining H3K4me3 occupancy upon RNF40 depletion. **C)** Differential Binding Analysis results showing H3K4me3 regulated (|log2FC|≥0.7, FDR<0.05) and unregulated regions (in purple). **D)** Heatmaps and respective aggregate profiles depicting changes of H3K4me3 occupancy in the identified gained (log2FC≥0.7, FDR<0.05, peak concentration≥;6.2), lost (log2FC≤-0.7, FDR<0.05, peak concentration≥6.2) or unregulated (|log2FC|≤0.2, FDR>0.1, peak concentration≥6.2) regions upon RNF40 depletion based on the DiffBind analysis results in C. **E)** Aggregate plots showing changes of H3K4me3 occupancy at TSS-associated regions of genes identified in RNA-seq analysis as robustly down-, up- (|log2FC|≥0.8, p-val<0.05) and unregulated (|log2FC|≤0.1, p-val>0.95) following RNF40 depletion. **F)** Quantification of changes in H3K4me3 peak width upon RNF40 depletion in regulated and unregulated genes. **G)** Left panel: classification of genes influenced by RNF40 depletion into Group A (simultaneous downregulation and H3K4me3 loss at TSS region), Group B (downregulation without H3K4me3 loss) and Group C (H3K4me3 loss at TSS region without expression changes). Right Panel: Group A genes were analyzed for pathway enrichment using the online EnrichR web tool (https://amp.pharm.mssm.edu/Enrichr3/). **H)** Box-Whiskers plot providing the median of normalized counts of the three gene groups (Group A, B and C). **I)** Changes in H3K4me3 occupancy at TSS-associated regions of group A, B and C genes. **J)** Aggregate plots of H2Bub1, H3K79me2, H3K36me3, H3K27ac, H3K9ac or RNApol II occupancy at TSS of group A, B and C genes in control HCC1954 cells (Accession number: GSE85158, GSE72956). All statistical tests: Mann-Whitney Test. **p-val<0.01, ***p-val<0.005.

**Fig. 6:**
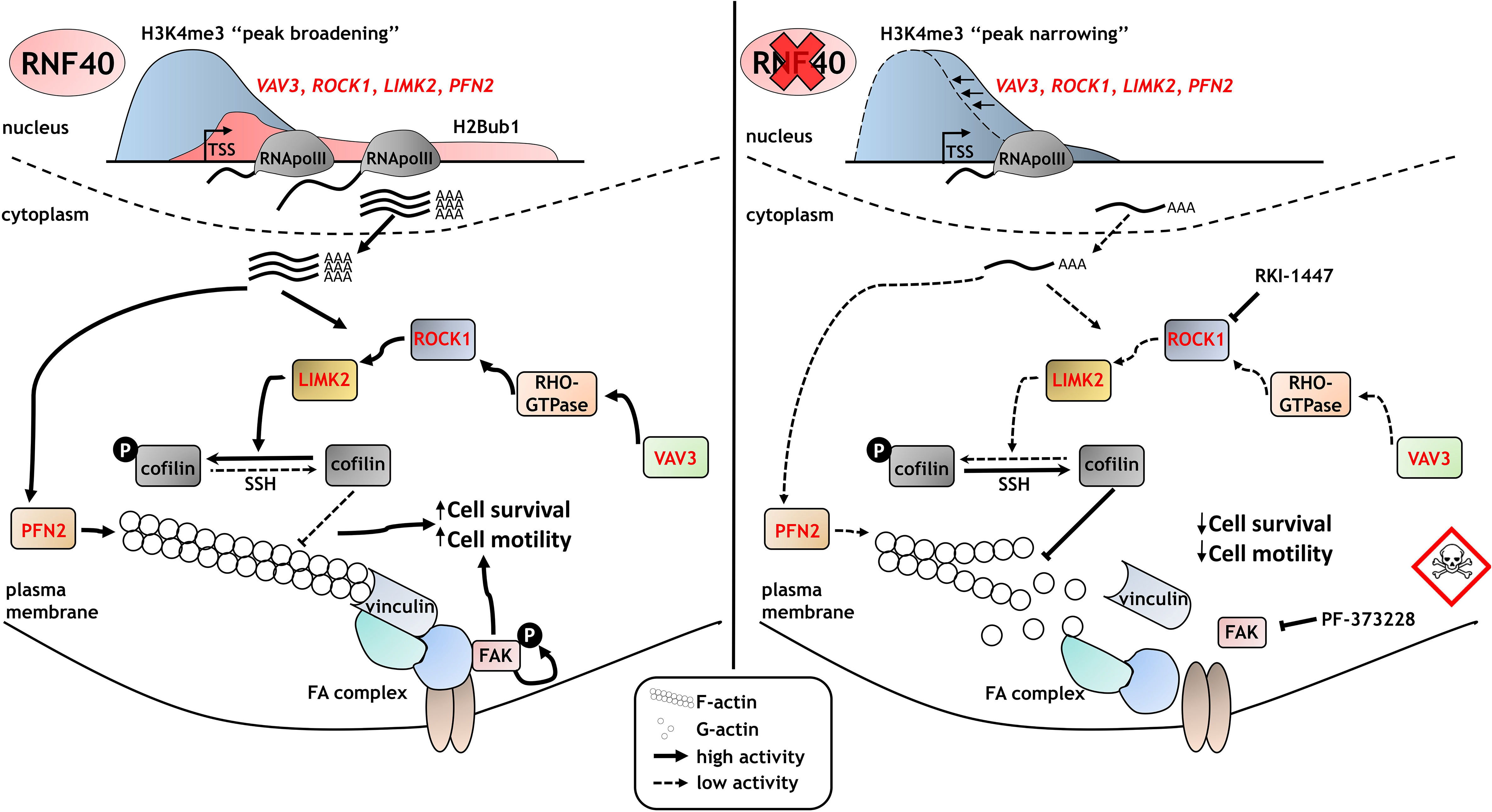
RNF40 enacts a tumor supportive role in HER2-driven mammary carcinoma via controlling the RHO/ROCK-dependent actin regulatory axis. RNF40-driven H2B monoubiquitination is essential for transcriptional activation of the RHO/ROCK/LIMK pathway components and for proper actin polymerization via a trans-histone crosstalk of histone 3 lysine 4 trimethylation (H3K4me3). Loss of RNF40 expression leads to the impairment of the H2Bub1-H3K4me3 axis, thereby dysregulating the actin dynamics and disrupting the focal adhesions (FA) and their pro-survival activity via FAK in HER2-positive breast cancer cells.

To further investigate the behavior of H3K4me3 occupancy at the TSS of regulated genes, we plotted H3K4me3 occupancy on robustly down-, up- and unregulated genes under control or RNF40-depleted conditions. Consistent with our analyses based on changes in H3K4me3 occupancy, genes downregulated upon RNF40 silencing displayed the most prominent decrease in H3K4me3 in the gene body (the 3’ end of the peak) compared to unregulated or upregulated genes (Fig. 5E-F). Importantly, a significant fraction of downregulated genes (162 out of 324) showed a concomitant decrease in H3K4me3 near the TSS (“Group A” in Fig. 5G). Moreover, Group A was enriched for genes involved in the actin dynamics regulatory pathway. The decrease in H3K4me3 spreading into the body of the *ROCK1, LIMK2* and *VAV3* genes could be validated by ChIP-qPCR (Fig. S5E). As a control, we identified a group of genes with a similar size and similar expression range as the Group A (Group C), whose expression was not affected by RNF40 knockdown, but was characterized by a milder reduction of H3K4me3 occupancy (Fig. 5G and Fig. 5H). Under normal culture conditions, genes of Group A harbored lower H2Bub1 levels than the genes of Group C, but comparable H3K4me3 levels and peak height (Fig. 5J and Fig. S5F-G). Interestingly, Group A genes presented a more profound H3K4me3 peak narrowing upon RNF40 depletion compared to the control group. The 162 genes found to be downregulated at the mRNA level but not showing any H3K4me3 loss (Group B) showed overall lower expression values and displayed only negligible levels of H2Bub1 across their gene body (Fig. 5H-J and Fig. S5G). We therefore concluded that the genes within Group B may likely be indirect downstream targets of RNF40-mediated H2B monoubiquitination. Together, these findings support the hypothesis that the actin regulatory gene network is dependent on direct epigenetic regulation by RNF40 through modulation of H2Bub1 and a trans-histone cross-talk with H3K4me3 levels in HER2-positive BC cells.

To further characterize the epigenetic differences distinguishing Group A and C, and which may help to explain the RNF40-dependency of Group A genes, we analyzed ChIP-seq data for several other histone modifications in HCC1954 cells (44,45). These analyses revealed that the occupancy of the active histone marks H3K27ac and H3K9ac was slightly higher at the TSS of Group C genes in comparison to Group A, while the elongation-associated modifications, H3K36me3 and H3K79me2, were dramatically higher in the gene body of Group C. Accordingly, RNA Polymerase II (RNApol II) occupancy was also higher on genes in Group C compared to the other groups (Fig. 5J). Together, when compared to the genes within Groups A and B, genes within Group C display a more pronounced occupancy of epigenetic marks associated with active gene transcription (46). Thus, these additional epigenetic modifications may help to compensate for the loss of H2Bub1 following RNF40 depletion, whereas lower levels of these active marks on Group A genes may render these to be more sensitive to changes in H2Bub1 occupancy.

In summary, we conclude that RNF40 is a major epigenetic regulator of the actin regulatory gene network in HER2-positive BC cells via H2B monoubiquitination and the downstream trans-histone control of H3K4me3 occupancy in the transcribed region.

## Discussion

H2B monoubiquitination has previously been reported to serve a tumor suppressive function with its levels gradually decreasing during cancer progression. Interestingly, the role of RNF20, a subunit of the obligate heterodimeric RNF20/RNF40 E3 ubiquitin ligase complex catalyzing the deposition of H2Bub1, is more contradictory and seems to exert opposing functions depending on cancer type or subtype (16,17). To date, only few studies focused on RNF40 expression in cancer. Upon examination of a cohort of both primary BC and brain metastases, we identified the loss of RNF40 expression and H2Bub1 as rare events in primary and metastatic HER2-positive lesions. Publically available datasets corroborate our results, showing only a very low rate of genetic alterations (<1%) causing loss of RNF40 function in BC (cbioportal.org, data not shown). Interestingly, the same datasets report a much higher frequency of *RNF40* locus amplification in malignancies of the breast (4-6%) accompanied by increased *RNF40* expression levels in tumors compared to normal tissues (TCGA dataset). Additionally, high expression levels of *RNF40* were associated with an unfavorable outcome in HER2-positive BC patients. Finally, the genetic model for RNF40 loss in endogenous HER2-driven mammary carcinomas used in this study supported the human patient data, arguing for a tumor-supporting role for RNF40 in HER2-dependent BC. Together, our data do not support a general tumor suppressive function of RNF40 and H2Bub1.

Upon investigating the transcriptional and molecular epigenetic mechanisms rendering HER2-positive BC cells critically dependent upon RNF40, we observed that loss of RNF40 had a profound impact on the deposition of the H3K4me3 histone mark leading to a significant “peak narrowing” in the transcribed region downstream of the TSS on regulated genes. The crosstalk between H2Bub1 and H3K4me3 has been intensively studied in the past and has been attributed to the trans-regulation of the histone methyl transferase activity of the COMPASS family of H3K4 methyltransferases by H2Bub1 (8,22,47). Our previous work revealed that RNF40 promotes the expression of a specific subset of genes displaying a high elongation rate via modulation of H3K4me3 peak broadening in a context-specific manner (22). Consistently, our new integrated datasets in HER2-positive BC not only further confirm that RNF40-dependent genes display a more profound tendency of H3K4me3 domain narrowing upon RNF40 depletion, but also show that these genes display a less pronounced accumulation of various activating epigenetic marks compared to RNF40-independent genes. Interestingly, RNF40-dependent genes also displayed lower H3K79me2 levels, another histone mark that was shown to function downstream of H2Bub1 to epigenetically regulate gene expression, implying that an additional epigenetic layer helps to control the transcriptional output of RNF40/H2Bub1-independent genes (28). We therefore hypothesized that this specific group of genes is rendered particularly sensitive to H2Bub1 loss upon RNF40 depletion due to their overall less active chromatin status.

Strikingly, a large fraction of the genes identified in this study as being RNF40-dependent are well known effectors of the actin regulatory pathway. Next to the reported implication of RNF40 in DNA damage response (48), replication stress (14), microtubule spindle organization (49), inflammation (18) and regulation of hormone receptor activity (12,20), the discovery that the maintenance of actin dynamics critically depends upon RNF40 in HER2-positive BC is both new and of significant interest. Notably, HER2-positive BC cells were previously shown to heavily rely on intact actin dynamics for cancer cell viability, motility and metastasis (39,50). Importantly, we specifically identified *VAV3, ROCK1, LIMK2* and *PFN2* as RNF40-dependent genes and confirmed the functional consequence of their impaired expression, which resulted in decreased cofilin phosphorylation both *in vitro* and *in vivo*, and decreased F-actin abundance and impaired actin dynamics. Importantly, we identified the ROCK1 kinase as a central RNF40-regulated factor controlling the actin regulatory pathway, as its inhibition via the potent ROCK inhibitor RKI-1447 was able to phenocopy the impaired tumorigenic phenotype caused by RNF40 loss. Interestingly, activation of the actin cytoskeleton signaling pathway by treating RNF40-depleted cells with an S1PR3 agonist partially rescued these effects. Therefore, these data strongly suggest that the imbalance in the control of actin dynamics in RNF40-depleted cells is largely a ROCK1-dependent phenomenon and support a previous study displaying the anti-tumorigenic effect of RKI-1447 in HER2-driven BC *in vivo* (37).

In addition to the central role of actin cytoskeleton dynamics in controlling cellular migration, the ROCK and focal adhesion kinase signaling pathway also has a critical function in suppressing apoptosis (37,51). While we previously identified a role for RNF40 in suppressing apoptosis in colorectal cancer cells via expression of anti-apoptotic members of the BCL2 family of proteins (19), our current results suggest that RNF40 suppresses programmed cell death in HER2-positive BC in a distinct manner via maintenance of ROCK-dependent focal adhesion kinase signaling. Indeed, focal adhesion structures decreased in size with a concomitant decrease in FAK kinase activity upon either RNF40 depletion or ROCK1 inhibition. Our data suggest that RNF20/RNF40-driven H2B monoubiquitination plays a decisive, context-specific function in HER2-positive BC by controlling the actin regulatory circuit and the downstream focal adhesion kinase-driven signaling cascade to maintain both anti-apoptotic signaling and control cellular migration in cancer cells. It is therefore attractive to speculate that a simultaneous inhibition of the RNF20/RNF40 E3 ubiquitin ligase activity together with inhibition of either ROCK1 or FAK might provide synergistic effects in the treatment of HER2-positive BC.

Together, our data support a context-dependent role of RNF40 and H2B monoubiquitination in breast carcinogenesis and suggest that the RNF20/RNF40 E3 ubiquitin ligase and/or its upstream regulators or downstream targets may serve as attractive targets for the development of new anti-cancer strategies in HER2-positive BC.

## Acknowledgments

We would like to thank S. Bolte, N. Molitor and the staff of the European Neuroscience Institute Göttingen for assistance in the animal handling administration, F. Alves (Translational Molecular Imaging, Max Planck Institute for Experimental Medicine, Göttingen) for access to the IncuCyte^®^ Live Cell Analysis System (Sartorius AG), S. Lutz (Institute of Pharmacology and Toxicology, University Medical Center Göttingen, Göttingen) for providing the vinculin antibody and M. Dobbelstein for reagents for measuring caspase 3/7 activity (Department of Molecular Oncology, Göttingen).

## Materials and Methods

### Animal handling and mouse model generation

Animals were housed under specific pathogen-free (SPF) conditions and in accordance with the animal rights laws and regulations of Lower-Saxony (LAVES, registration number #15/1754). For more details, please refer to the Supplementary Data.

### Histology of human and murine tissues and publically available dataset analyses

Tissue microarrays of human primary and metastatic breast cancer were generated at the University Medical Center Hamburg Eppendorf Germany (local ethical committee approval number: OB/V/03 and MC-267/13, respectively) in accordance with the ethical standards of the 1964 Declaration of Helsinki. RNF40 and H2Bub1 scoring was established based on the staining intensity (null=no detectable staining, low=weak staining intensity, high=strong staining intensity). Detailed staining procedures, antibodies used for immunohistochemical staining are provided in the Supplementary Data.

### Publically available datasets

The Kaplan-Meier plotter (kmplot.com) and The Cancer Genome Atlas (TCGA)-derived publically available datasets were used to examine the association of *RNF40* expression with Relapse-Free Survival (RFS) or Overall Survival (OS) of HER2-positive BC patients. Please refer to the Supplementary Data for BC subtype classification parameters. Publically available datasets for histone modifications of active transcription in HCC1954 cells (GSE85158 and GSE72956) were downloaded from Gene Expression Omnibus (www.ncbi.nlm.nih.gov/geo/) (44,45).

### Cell culture, transfections and functional assays

HCC1954 (ATCC^®^ CRL-2338™) and SKBR3 (ATCC^®^ HTB-30™) cells were purchased from the American Type Culture Collection (ATCC). siRNA transfections were performed using Lipofectamine^®^ RNAiMAX (Invitrogen) according to the manufacturer’s guidelines. Proliferation kinetics as well tumor spheres were recorded using Celigo^®^ S imaging cytometer (Nexcelom Bioscience LLC) and IncuCyte^®^ Live Cell Analysis System (Sartorius AG). Colonies and migrated cells from trans-well assay were washed with PBS, fixed, stained and scanned with an Epson Perfection V700 Photo. Detailed protocols for siRNA transfection of both cell lines are available in the Supplementary Data.

### Immunofluorescence microscopy

Cells were plated and transfected on coverslips and grown for another 72 h. Cells were then washed with PBS, cross-linked with 4% paraformaldehyde and permeabilized with 1% Triton X-100 in PBS or TBS for 10 min, blocked for 1 h and incubated with the primary antibody overnight. Coverslips were washed and secondary antibody was applied with DAPI and eventually Alexa555-phalloidin for 1 hour at room temperature. Coverslips were washed and mounted on microscope slides. A detailed protocol as well for a list of antibodies is available in the Supplementary Data.

### Microscopy

Immunohistochemistry (IHC) pictures were taken with a Zeiss Axio Scope A1. Bright-field images of cultured cells were taken with a Nikon Eclipse S100 inverted microscope and immunofluorescence pictures with a Zeiss LSM 510 Meta confocal microscope. Fluorescence intensity was quantified using the ImageJ software. Image analysis workflow is described in the Supplementary Data.

### Annexin and caspase 3/7 activity assay

For annexin V staining, cells were tryprinized and resuspended in binding buffer at 72 hours post-transfection and incubated with Annexin V-FITC (Southern Biotech) and propidium iodide (Sigma Aldrich) for 15 min at room temperature. Samples were analyzed using a Guava EasyCyte Plus flow cytometer (Guava Technologies).

The kinetic apoptosis assay using caspase 3/7 was performed according to the manufacturer’s instructions (CS1-V0002(3)-1, ViaStainTM Live Caspase 3/7 Detection Kit, Nexcelom). Scanning was performed at time points 24, 48 and 72 hours post transfection using a Celigo^®^ S imaging cytometer (Nexcelom Bioscience LLC). For detailed protocols, please refer to the Supplementary Data.

### ChIP library preparation and data analysis

Chromatin immunoprecipitation was performed as described previously (52) 72 hours after transfection with control or RNF40 siRNAs using antibodies against H2Bub1 (Cat. No. 5546S, Cell Signaling Technology) and H3K4me3 (Cat. No. C15410003-50, Diagenode). Next generation sequencing libraries were prepared using the Microplex Library Preparation kit v2 (Diagenode, Cat.No. C05010011) according to manufacturer’s instructions and samples were sequenced (single-end 50 bp) on a HiSeq4000 (Illumina) at the Transcriptome and Genome Analysis Laboratory (TAL) at the University Medical Center Göttingen. Processing of sequencing data was performed in the Galaxy environment provided by the “Gesellschaft für wissenschaftliche Datenverarbeitung mbH Göttingen” (galaxy.gwdg.de). Briefly, ChIP-seq reads were mapped to the hg19 reference genome assembly using Bowtie2 (version 2.3.2.2). PCR duplicates were removed using the RmDup tool (version 2.0.1). The deeptools suite (version 3.2.0.0.1) was utilized to generate normalized coverage files (bamCoverage), call peak changes (bigwigCompare), and to generate aggregate plots and heatmaps (computeMatrix and plotHeatmap). Occupancy profiles were visualized using the Integrative Genomics Viewer (IGV 2.4.8). Detailed analysis workflow is available in Supplementary Data.

### RNA library preparation and data analysis

RNA sequencing libraries were generated from HCC1954 cells at 72 hours post-transfection with the NEXTFLEX^^®^^ Rapid Directional RNA-Seq Kit (Bioo Scientific, Catalog #NOVA-5138-07) according to the manufacturer’s instructions and samples were sequenced (single-end 50 bp) on a HiSeq4000 (Illumina) at the TAL. RNA-seq data were processed in the Galaxy environment. Raw reads were trimmed (FASTQ Trimmer), mapped to the reference genome hg19 using TopHat (version 2.1.1) and read counts per gene was calculated with featureCounts. Finally, differential gene expression analysis and normalized counts were obtained using DESeq2. Detailed analysis workflow is available in Supplementary Data.

## Notes

This work was supported by funding from the Deutsche Krebshilfe to S.A.J. (111600), and to H.W. and K.P. (Priority Program “Translational Oncology”; 70112507).

**Conflict of interest:** F.W., E.P., A.P., R.L.K., K.P., H.W., M.G. and S.A.J. declare no conflict of interest.

